# Sex-specific molecular specialization and activity rhythm dependent gene expression changes in honey bee antennae

**DOI:** 10.1101/755728

**Authors:** Rikesh Jain, Axel Brockmann

## Abstract

Eusocial insects, like honey bees, which show an elaborate division of labor involving morphologically and physiologically specialized phenotypes provide a unique toolkit to study molecular underpinnings of behavior as well as neural processing. In this study, we performed an extensive RNA-seq based comparison of gene expression levels in the antennae of honey bee drones and foragers collected at different time of days and activity states to identify molecules involved in peripheral olfactory processing and provide insights into distinct strategies in sensory processing. First, honey bee drone and worker antennae differ in the number of olfactory receptor genes (ORs) showing a biased expression pattern. Only 19 Ors were higher expressed in drone antennae, whereas 54 Ors were higher expressed in workers. Second, drone antennae showed predominant higher expression of genes involved in energy metabolism, and worker antennae showed a higher expression of genes involved in neuronal communication. Third, drones and afternoon-trained foragers showed similar daily changes in the expression of major clock genes, *per* and *cry2*. Most of the other genes showing changes with the onset of daily activity were specific to drones and foragers suggesting sex-specific circadian changes in antennae. Drone antennae are specialized to detect small amounts of queen’s pheromone and quickly respond to changes in pheromone concentration involving energetically costly action potentials, whereas forager antennae are predominantly involved in behavioral context dependent detection and discrimination of complex odor mixtures which requires mechanisms of sensory filtering and neural plasticity.

## INTRODUCTION

In honey bees, males (drones) and females (workers and queens) exhibit a strong sexual dimorphism in the peripheral olfactory sensory system. Drone antennae have about seven times more olfactory poreplate sensilla and are specialized to detect even minute amounts of the queen’s sex-pheromone (Brockmann et al., 1998; Brockmann et al., 2006; Esslen and Kaissling, 1976; Wanner et al., 2007). In contrast, workers have a more generalized olfactory system with different, maybe broader, odorant response profile. For example, worker antennae house hair-like sensilla (S. basiconicum) that are absent on drone antennae, and worker antennal lobes exhibit about 170 isomorphic olfactory glomeruli, whereas drones have only 100 normal-sized and 4 macro glomeruli (Arnold et al., 1985; Brockmann and Brückner, 2001; Esslen and Kaissling, 1976; Flanagan and Mercer, 1989; Galizia et al., 1999; Kropf et al., 2014; Sandoz, 2006). These differences in the olfactory sensory systems correlate with the different behavioral functions. Drones have to find and mate with one queen at the same time outcompeting other drones (Brockmann et al., 2006; Gary, 1962; Koeniger et al., 2005; Ruttner, 1985). In contrast, workers do all the tasks needed to maintain the colony and organize the underlying division of labor; and as foragers, they have to learn and memorize odor mixtures that indicate different rewarding flowers (Frisch, 1967; Frisch and Aschoff, 1987). These different behavioral contexts suggest that the drone and worker antennal sensory systems may exhibit different sensory processing strategies and molecular adaptations (Burger et al., 2013; Grabe and Sachse, 2018). Wanner et al., (2007) already reported differences in the expression of olfactory receptor genes (Or) between drones and workers and showed that one of the male-biased expressed Ors (*Or11*) binds 9-ODA, the major sex-pheromone compound.

In this RNA-seq study, we first compared the antennal transcriptomes of drones and foragers, to identify gene expression that might reflect differences between a specialist and a generalist peripheral olfactory system involving innate and learned sensory processing (Amin and Lin, 2019; De Bruyne and Baker, 2008; Renou, 2014). In a second set of experiments we compared daily changes in antennal gene expression between drones, that only leave the hive for mating flights in the early afternoon, and foragers that were either entrained to visit a feeder in the morning or in the afternoon. This comparison allows to explore to which extent drone and worker antennae show similar or different daily changes in gene expression. One hypothesis would be that the daily gene expression changes might correlate with the molecular specialization of the two types of antennae. Finally, we expect to identify genes that are not directly involved in odorant detection, but likely play an important role in peripheral olfactory processing in insect antennae. Furthermore, these genes might indicate differences in the sensory processing in drone and worker antennae.

## MATERIALS AND METHODS

### Animals

In all experiments we used *Apis mellifera* colonies of naturally mated queens which consisted of about 8000 workers (i.e. 8 frames with approximately 1000 workers) and hundreds of drones. Colonies were acquired from a local beekeeper and maintained on the campus of the National Centre for Biological Sciences (NCBS), Bangalore, India.

### Daily drone flight activity

Daily drone flight activity was determined for three colonies on three different days during a period of two weeks (Oct 28, Nov 03 and Nov 10, 2017). On the experimental days numbers of drones leaving the hive entrance were counted every half an hour for 10 minutes from 7:00 to 19:00 hours (h). During this time of the year, sunset is at around 18:00 h in Bangalore. On these days, we also recorded temperature and humidity changes every minute using a data logger (EQ-172, Equinox, Valli Aqua And Process Instruments, Chennai, India).

### Collection of drones for antennal RNA-seq and qPCR

During daily mating flight activity, drones were caught at the hive entrance and color marked on the thorax. On the next day color-marked drones were collected at two different time points: 9:00 (inactive) and 14:00 h (active/mating flight time, also see Naeger and Robinson, 2016) from 3 different colonies (5 bees per time point per colony). At 9:00 h drones were collected from inside the colonies and at 14:00 h they were collected from the entrance before they started the mating flights. In a separate experiment we collected color-marked drones from one of the three colonies at 6 different time points: 6:00, 10:00, 14:00, 18:00, 22:00 and 2:00 (10 bees per time point) to determine daily expression changes of four major clock genes i.e. *period* (*per*), *cryptochrome2* (*cry2*), *cycle* (*cyc*) and *clock* (*clk*). Night collections were done using dim red light. All collected drones were immediately flash frozen in liquid nitrogen.

### Collection of time-trained foragers for antennal RNA-seq

An *A. mellifera* colony was transferred in an enclosed outdoor flight cage to entrain the foraging activity of the workers to a distinct time of the day. First, the colony was allowed to adjust to the new environment for 10 days. During this period the sugar and pollen feeders were presented for the whole day. The sugar feeder was a yellow plastic plate surrounded with 4 filter papers containing a 5µl drop of 100 times diluted linalool racemic mixture (Sigma-Aldrich, St. Louis, Missouri). Then, for the time-training, the sucrose reward (1M sucrose solution) was presented either from 8:00 to 10:00 h (morning training) or from 13:00 to 15:00 h (afternoon training) for 10 consecutive days. Time for the afternoon training was chosen according to the drone flight time. Two different colonies were used for morning and afternoon training. Every day after the training time the feeder was cleaned with ethanol and linalool scented filter papers were replaced with fresh unscented filter papers. This cleaned empty feeder was available for the remaining time of the day. On the 8^th^, 9^th^ and 10^th^ day of training, foragers visiting the feeder were marked on their thorax with different colors, one type of color each day, to identify the frequently visiting foragers. On the 11th day, the feeder was not presented and the foragers that had all 3 color marks were collected at 9:00 and 14:00 h from both the colonies (10 bees per time point per colony). Collected foragers were immediately flash frozen in liquid nitrogen.

### RNA isolation from the antennae

Collected honey bee samples were transferred from liquid nitrogen onto dry ice and the entire antennae (i.e. scape, pedicel and flagellum) were cut off. We pooled 10 antennae from 5 bees for RNA-seq samples and 4 antennae of 2 bees for qPCR samples. Total RNA was isolated using Trizol® method (Invitrogen, Carlsbad, CA). Samples were treated with DNaseI (Invitrogen, Carlsbad, CA) for 10 minutes to remove any possible DNA contamination. Final RNA concentration was measured using nanodrop and quality was confirmed by running an agarose gel.

### qPCR

0.5µg-1µg of total RNA was converted into cDNA using Superscript III and oligo d(T)16 primers (Invitrogen, Carlsbad, CA). QPCR was performed using KAPA SYBR FAST qPCR Master Mix (Kapa Biosystems, Wilmington, MA) in 7900HT Fast Real-Time PCR system (Applied Biosystems, Carlsbad, CA). Triplicate reactions (10ul reaction mix) for all the biological replicates of all 6 time points samples (n=5 per time point) were run in parallel on the same 384-well plate. This restricted us to analyze just one of the clock genes (S1 Table) and *ribosomal protein49* (*rp49*) (an internal control gene) (Jain and Brockmann, 2018) per plate. We also ran standard curve for both primers on the same plate using a separate stock cDNA. Final gene expression calculation was based on the linear values interpolated from the standard curves. Efficiency of all the primers were between 95-100%. QPCR reactions with bad dissociation curve were discarded from the analysis.

### RNA-seq

Antennal transcriptomes of drones (n=3 per time point), morning-trained foragers (n=2 per time point) and afternoon-trained foragers (n=2 per time point) were sequenced at 2 different time points (9:00 h and 14:00 h). Total RNA was shipped on dry ice to AgriGenome Labs (Kochi, India). RNA quality was further checked on Agilent Tapestation and Qubit. Libraries were prepared using TruseqRibozero gold + Truseq mRNA stranded library prep Kit. Sequencing was performed on an Illumina NextSeq500 platform and around 120 millions of 75-bp-long paired-end reads were generated.

### Data analysis

#### qPCR

We used cosinor package (Mutak, 2017) in R (R Core Team, 2017) to fit a 24 hour cosine model {y = intercept + amplitude * cos(2*pi(x - acrophase)/24)}(Nagari et al., 2017) in the circadian genes expression data. We performed a non-parametric JTK cycle analysis (Hughes et al., 2010; Patton et al., 2014) to detect daily rhythmicity in clock genes expression and Kruskal-Wallis test to show the differences in mRNA levels with time of the day (Table 1).

**Table 1.** Non-parametric JTK cycle analysis and Kruskal-Wallis test for qPCR data.

| Gene Name | LAG | AMP | ADJ. P | p-value<br>(Kruskal-Wallis test) |
| --- | --- | --- | --- | --- |
| *Cry2* | 4 | 6.62 | 7.96E-10 | 1.31E-04 |
| *Per* | 4 | 3.19 | 2.02E-08 | 2.71E-04 |
| *Cyc* | 10 | 0.53 | 3.54E-04 | 4.49E-03 |
| *Clk* | 6 | 0.01 | 1 | 0.702 |

#### RNA-seq

Approximately 7-10 million pairs of 75-bp-long reads per sample were mapped to *Apis mellifera* genomeHAv3.1 (Wallberg et al., 2019) using STAR (Dobin et al., 2013). The alignment rate was more than 75% (75.15% to 86.82%) for all the samples. The number of reads aligning to each gene were counted using featureCounts (Liao et al., 2014). The differentially expressed genes (DEGs) with p-adj. less than 0.05 (Wald test) were identified using DESeq2 (Love et al., 2014). Pathview package (Luo and Brouwer, 2013) in R was used to integrate the DEGs data to relevant pathway graphs from KEGG and to visualize. In addition, GAGE package (Luo et al., 2009) in R was used for geneset enrichment analysis (GSEA) using normalized count data from featureCounts and the KEGG pathway database (Kanehisa and Goto, 2000). Gene Ontology (GO) enrichment analysis was done using g:Profiler (Raudvere et al., 2019) keeping alpha of 0.05 as cut off for significance.

## RESULTS

### 1. Drones and worker antennae show sex-specific molecular specialization indicating different sensory processing strategies

Drone and worker antennae showed distinct and prominent transcriptomic profiles. The 14 RNA-seq samples separated in 2 clear sex-specific clusters (Fig 1), and we could identify 3998 differentially expressed genes (DEGs) (*p.adj*<0.05) between drones and foragers (S1 Table). Out of these 3998 genes 1815 were higher expressed in drones and 2183 were higher expressed in foragers (Fig 1).

**Fig 1.**
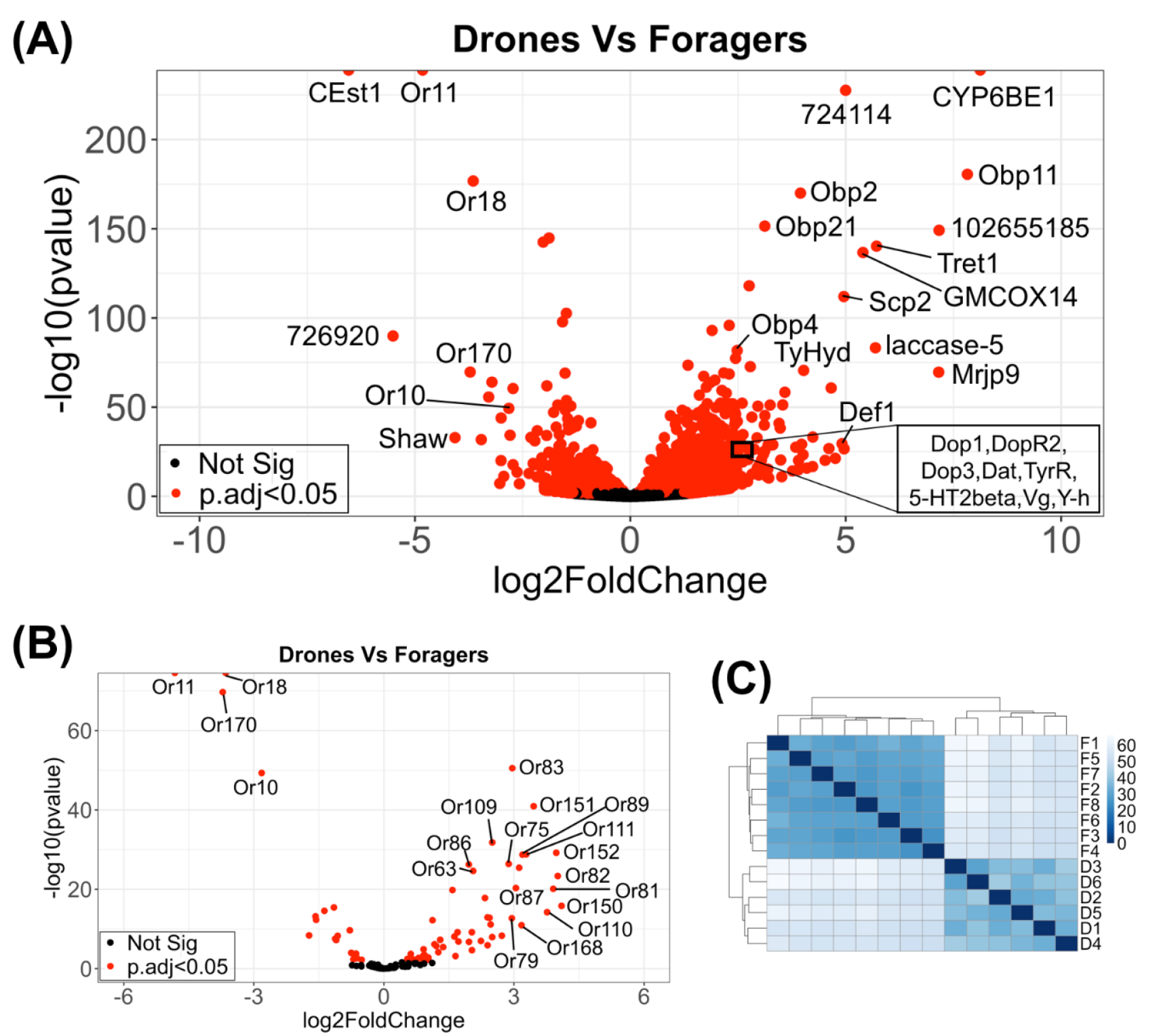
Sexual differences in antennal transcriptome. (A) Differential expression (DESeq2) of all the genes across drone versus forager antennae. (B) Differential expression (DESeq2) of odorant receptors (Ors) genes across drone versus forager antennae. (C) Sample distance heatmap based on variance stabilized RNA-seq read-count from DESeq2. Drone samples D1, D2 and D3 are collected in the morning and D4, D5 and D6 are collected in the afternoon. Similarly, forager samples F1, F2, F3 and F4 are collected in the morning and F5, F6, F7 and F8 are collected in the afternoon. Forager F1, F2, F5 and F6 are the afternoon-trained foragers and remaining are morning-trained foragers.

Remarkably, among the DEGs with the highest expression differences in our study were almost all the genes that previously were reported to be differently expressed: *Or10*, *Or11* (the 9-ODA olfactory receptor), *Or18*, *Or170* and *carboxyl esterase1* (*CEst1*) were higher expressed in drone antennae (Wanner et al., 2007); and *Or63*, *Or81*, *Or109*, *Or150*, *Or151*, *Or152*, *Obp2*, *Obp4*, *Obp11*, *Obp16*, *Obp19*, *Obp21*, *CSP6* and *Cyp6BE1* were higher expressed in worker antennae (Fig 1A and S1 Table) (Forêt and Maleszka, 2006; Wanner et al., 2007). However, in contrast to these studies, our RNA-seq analysis identified total 73 Ors genes showing significant expression differences (*p.adj*<0.05) between drone and forager antenna, whereof 19 (12 with log2fold change>1) were higher expressed in drones and 54 (40 with log2fold change>1) higher in foragers (Fig 1B and S2 Table). In addition to the olfactory receptor genes we found a higher expression of *Obp1*, *ionotropic receptor 21a* and *the gustatory receptor for sugar taste 43a* (=*Amgr3*) in drone antennae, and higher expression of the*, Obp12, and CSP3* in forager antennae.

Besides the genes that are obviously involved in olfaction we found a number of significantly different genes (log2fold change>1, *p.adj*<0.05) that either could be involved in olfactory sensory processing or other important function of the antennae (Table S1). Drone antennae showed a higher expression of the two major sex-determining genes, *complementary sex determiner* (*csd*) and *feminizer* (*fem*), the voltage-gated potassium channel (*Shaw* = Shaker cognate w), *nitric oxide synthase* (*NOS*), and *glutathione S-transferase* (*GstD1*, *GstS4*) the latter three likely playing a role in odorant detection, olfactory transduction and cellular signaling.

Most interestingly, forager antennae showed a higher expression of several biogenic amine receptors: *Dop1*, *Dop3*, *DopR2* (dopamine receptors), *5-HT2alpha*, *5-HT2beta* (serotonin receptors), *TyrR* (tyramine receptor), several glutamate receptors (*ionotropic glutamate receptor*, *glutamate receptor 1* and *vesicular glutamate transporter 1*), enzymes of the tyrosine/dopamine biosynthesis pathway (*tyrosine hydroxylase* and *tyrosine aminotransferase*), as well as several genes involved in neuropeptide signaling (*adipokinetic hormone receptor*, *tachykinin*, *prohormone-2* and*neprilysin-4*). Further, theTRPV channel *nanchung* showed a higher expression in forager antennae. *Nanchung* was reported to be expressed in the Johnston’s organ and involved in hearing and gravity perception (Sun et al., 2009). Mondet et al (2015) previously showed a higher expression of *nanchung* in forager antennae compared to nurse antennae. Finally, *vitellogenin* (*Vg*), several genes of the *major royal jelly proteins* and *the yellow proteins*, all likely be involved in sex- and caste-specific behaviors (*Mrjp1*, *Mrjp3*, *Mrjp8*, *Mrjp9,Y-h*, *Y-y*, *Y-e3*, *Y-f*) as well as the immune genes *defensin* (*Def1*, *Def2*), *abaecin*, *apidaecin1* (*Apid1*), and *transferrin1* (*Tsf1*) were higher expressed in worker antennae.

Gene set enrichment analysis using KEGG pathway database revealed significant (q-value<0.1) upregulation of 65 biological pathways in drone and 4 biological pathways in forager antennae (S3 Table). Two of the most significant pathways (lowest q-values) in drone antennae were oxidative phosphorylation (ame00190) (Fig2A) and protein processing in endoplasmic reticulum (ame04141) (Fig 2B). In contrast, in worker antennae, ligand-receptor interaction (ame04080) (Fig 3A) and tyrosine metabolism (ame00350) (Fig 3B) were the most significant.

**Fig 2.**
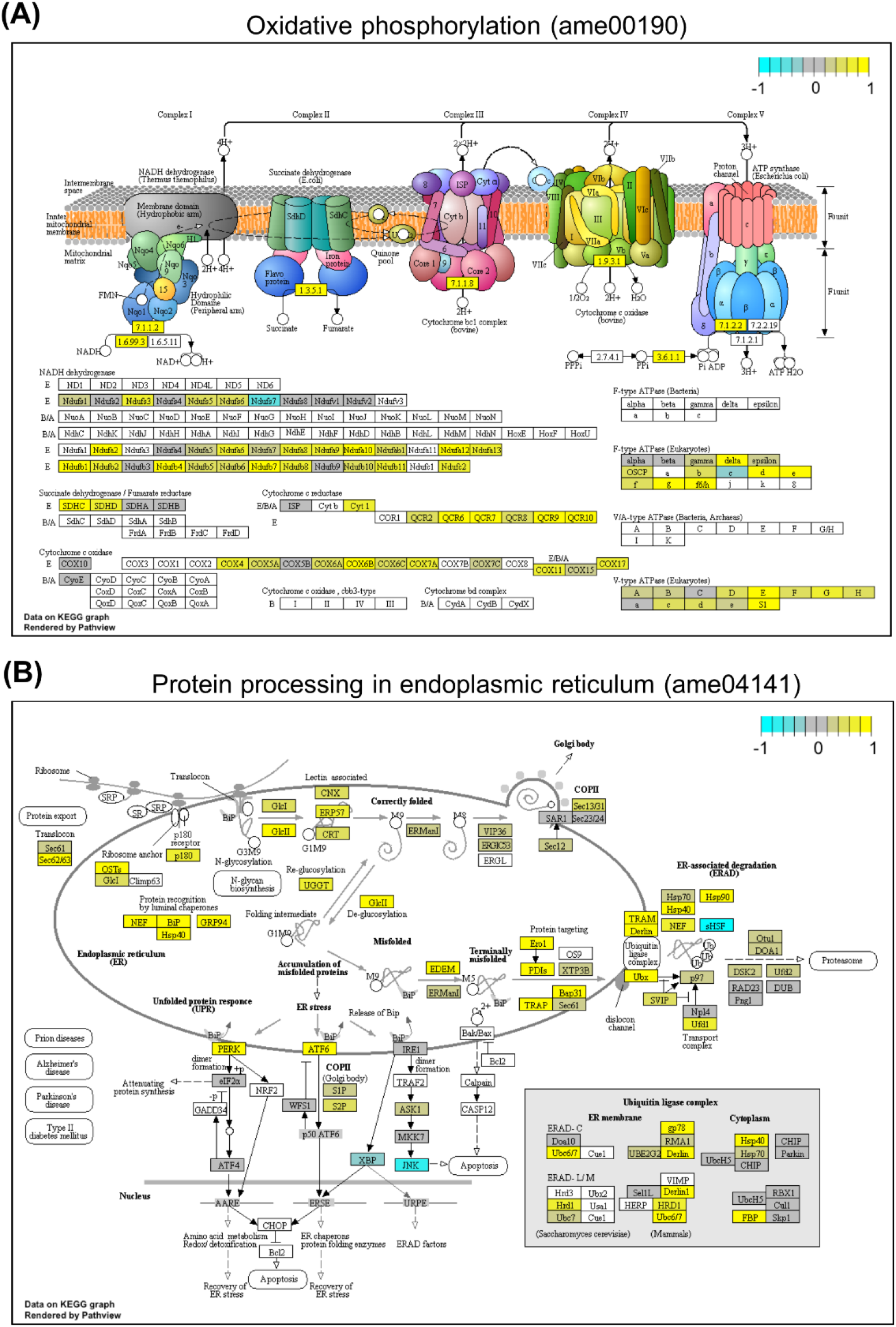
Top two significantly enriched KEGG pathways in drone antennae. Gene set enrichment analysis was performed using gage package and gene expression data was integrated to relevant KEGG pathways using pathview package in R. Yellow (positive values) highlighted genes are higher expressed in drones, cyan (negative values) highlighted genes are lower expressed in drones or higher expressed in foragers. Genes with gray background do not show expression differences between drones and foragers. Genes with transparent background are not found or annotated in honey bees.

**Fig 3.**
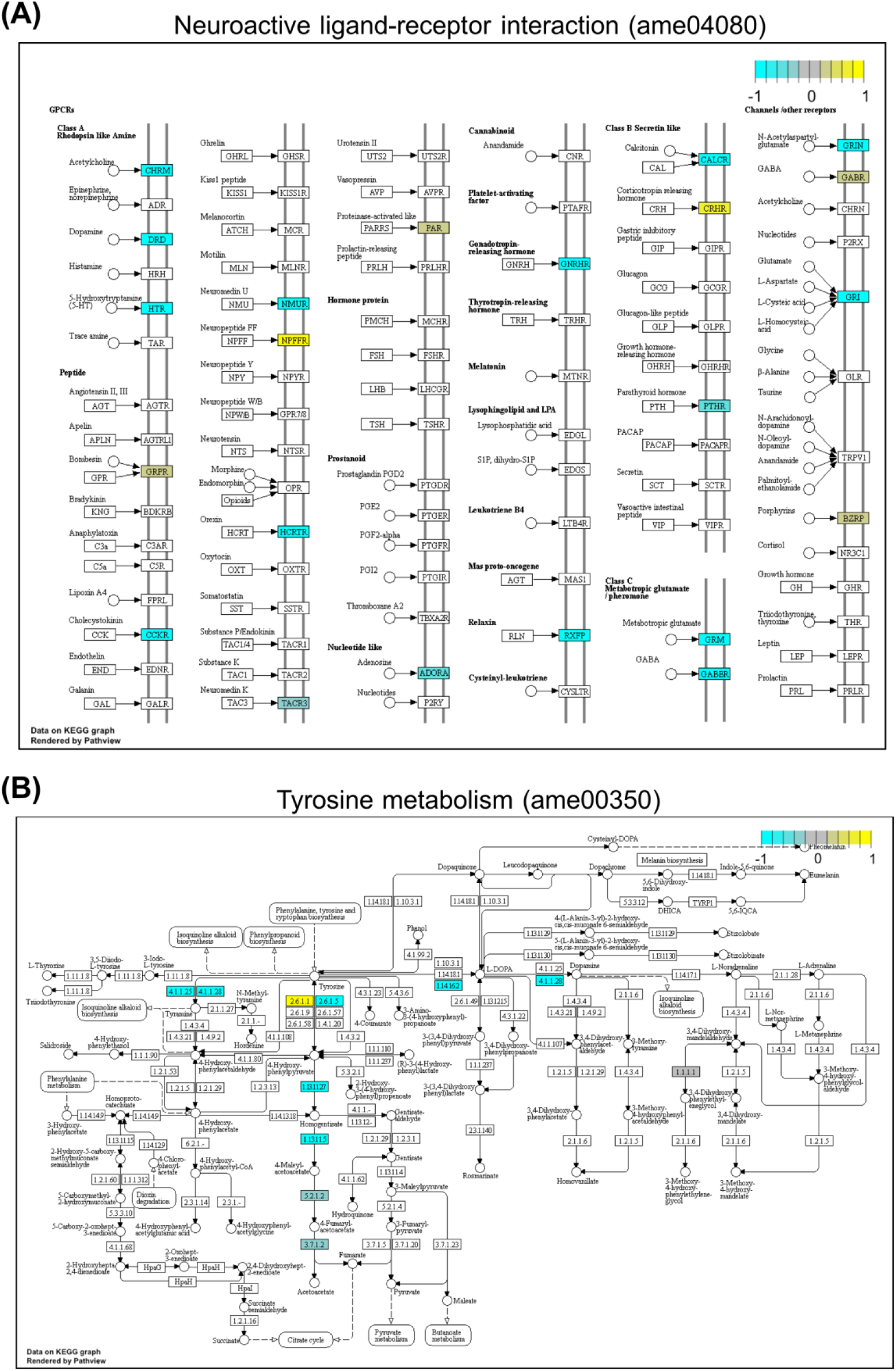
Top two significantly enriched KEGG pathways in foragers antennae. Gene set enrichment analysis was performed using gage package and gene expression data was integrated to relevant KEGG pathways using pathview package in R. Cyan (negative values) highlighted genes are higher expressed in foragers and yellow (positive values) highlighted genes are lower expressed in foragers or higher expressed in drones. Genes with gray background do not show expression differences between drones and foragers. Genes with transparent background are not found or annotated in honey bees.

Gene Ontology (GO) enrichment analysis using DEGs with more than 2 fold expression differences (471 DEGs in drones and 914 in foragers) showed significant enrichment of 53 and 105 GO terms in drones and foragers respectively (*p*<0.05; S4 Table). Significantly enriched GO terms in drones include catalytic activity (GO:0003824), odorant binding (GO:0005549), metabolic process (GO:0008152), protein folding (GO:0006457), cytoplasmic part (GO:0044444) and mitochondria (GO:0005739). In foragers, some of the significantly enriched GO categories were signaling receptor activity (GO:0038023), molecular transducer activity (GO:0060089), regulation of cellular process (GO:0050794), signaling (GO:0023052), sensory perception (GO:0007600), integral component of membrane (GO:0016021) and extracellular region (GO:0005576).

### 2. Antennae of drones performing afternoon mating flights and antennae of foragers entrained to forage in the afternoon show similar clock gene expression patterns

Drones of three *A. mellifera* colonies maintained at NCBS campus performed mating flights between 13:00 and 15:00 hours on all three observation days (Fig 4). Flight activity did not differ among colonies and experimental days. During these two hours the temperature was about 25°C and the relative humidity was around 60-70%.

**Fig 4.**
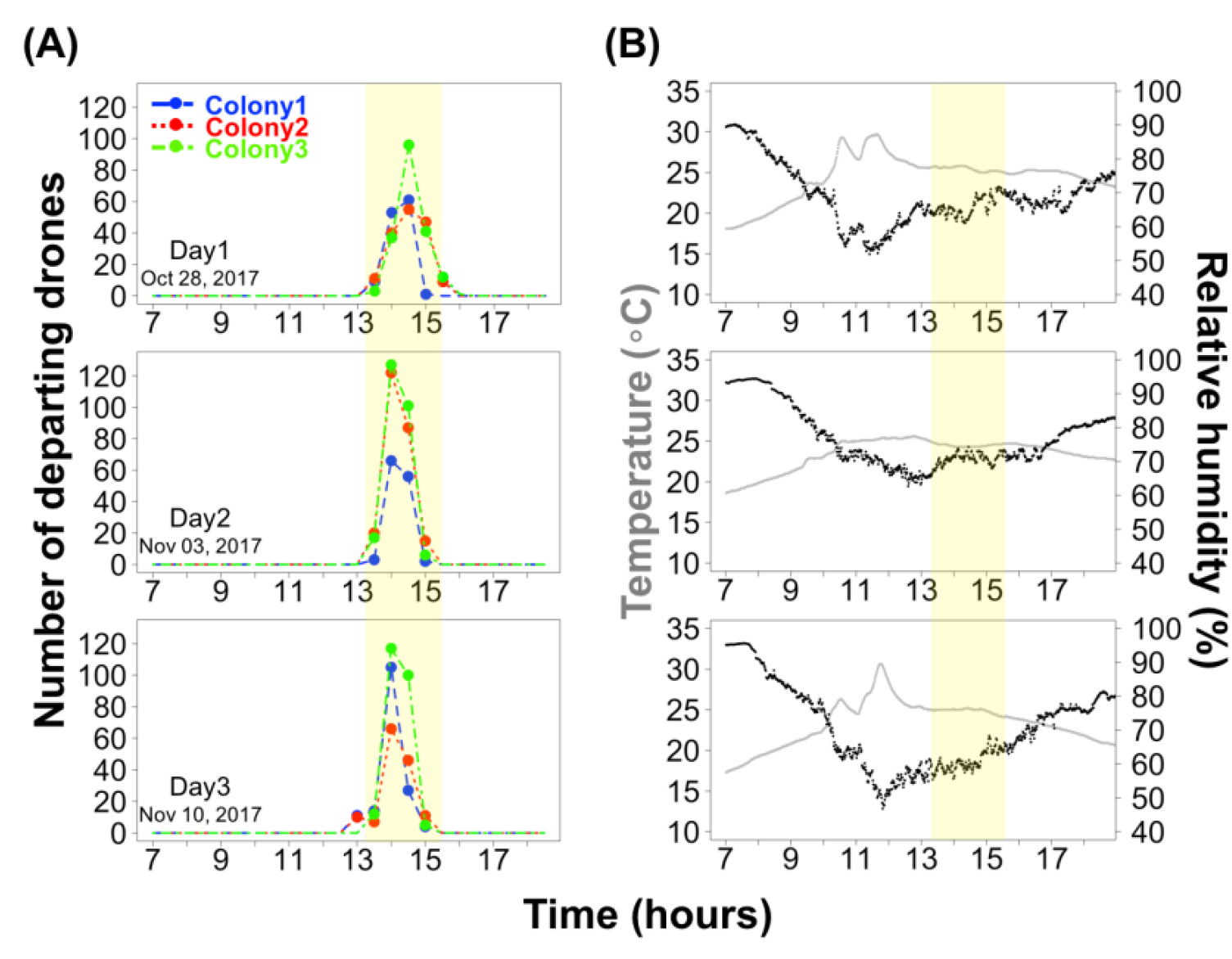
Drone flight timing of *A. mellifera* in Bangalore. (A) Number of departing drones, counted for the first 10 minutes every half an hour over 3 days from 3 different colonies (color code). (B) Temperature and humidity was recorded at each minute on all 3 days.

The antennae of drones performing mating flights showed significant 24-hour daily rhythms in the mRNA levels of major clock genes (n=5 per time point; Fig 5 and Table 1). *Cry2* and *per* mRNA levels peaked during early morning, while the *cyc* mRNA level was highest during the afternoon. *Clk* mRNA levels did not change. *Cry2* oscillated with higher amplitude than *per* and *cyc*. This expression pattern of clock genes is similar to that of afternoon-trained foragers (Jain and Brockmann, 2018; Spangler, 1972).

**Fig 5.**
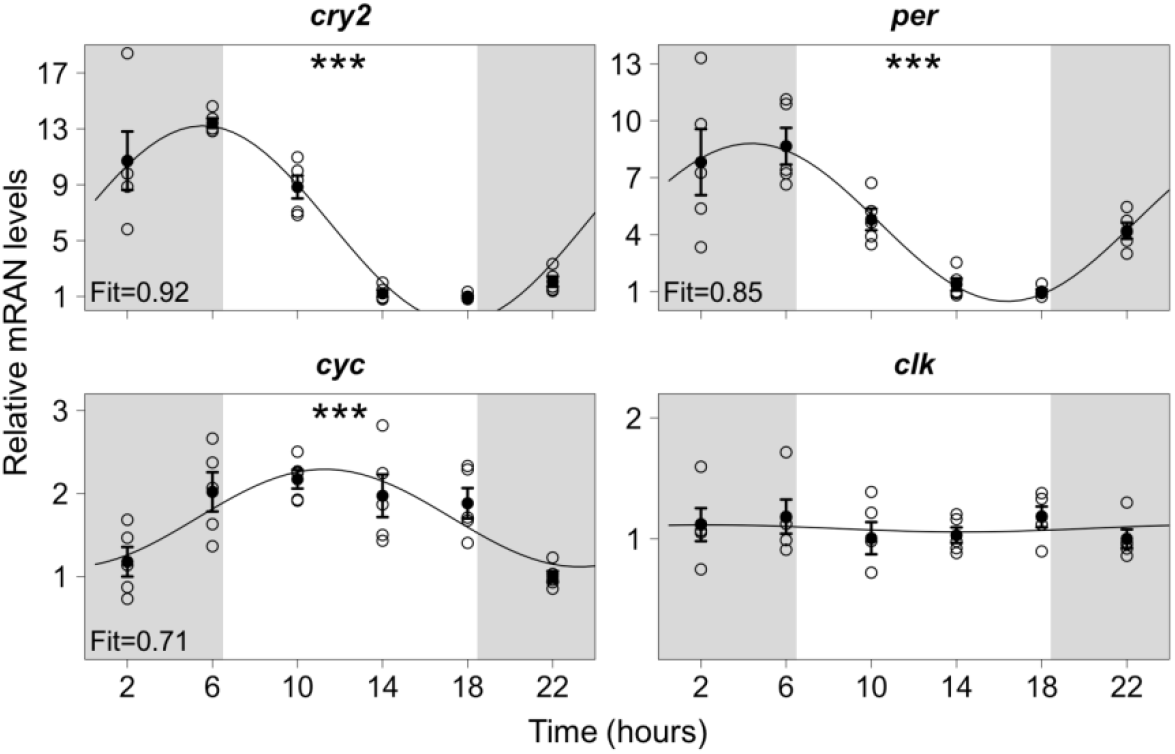
Clock genes mRNA rhythm in drone antennae. Open circles indicate individual qPCR measurements from 4 pooled antennae (n=5 per time point), normalized against the internal reference gene, *rp49*. Filled circle represents the mean ± SEM. Curved lines through the data points correspond to the best fitted 24-hour cosine models. Fit values for the cosine models are indicated at the bottom left of the plots. Statistical significance of daily mRNA rhythms are presented with asterisks (\*\*\**p*<0.005, Kruskal-Wallis test and JTK cycle analysis) at the top center of each plot. Gray shades in the background of each plot indicate the night time i.e. 18:30 to 6:30.

Column named LAG, AMP and ADJ.P are from JTK cycle analysis (Hughes et al., 2010). LAG represents approximate time of the day (in hours) at which the gene expression cycle reaches its maximum and AMP stands for amplitude of the gene expression cycle (relative mRNA levels in arbitrary units). ADJ.P is for adjusted p-values.

### 3. Daily activity rhythm affects antennal gene expression in drones and foragers

We found 75, 72 and 70 DEGs (*p.adj*< 0.05) between morning (9:00 h) and afternoon (14:00 h) in the antennae of drones, foragers visiting a feeder in the morning (morning-trained foragers), and foragers trained to visit a feeder in the afternoon (afternoon-trained foragers), respectively (S5 Table).

Two DEGs, *per* and *heat shock protein 90* (*hsp90*), showed change in expression between morning and afternoon in all 3 groups compared (Fig 6A and 6B). Similar to our qPCR results, expression of *per* was strongly associated with the daily activity rhythm. In drones and afternoon-trained foragers *per* mRNA level was higher in the morning and in morning-trained foragers *per* expression was higher in the afternoon. In contrast, the expression change of *hsp90* was sex-specific. In drones *hsp90* was higher expressed in the afternoon whereas in foragers it was higher expressed in the morning independent of the activity rhythm of the forager group.

**Fig 6.**
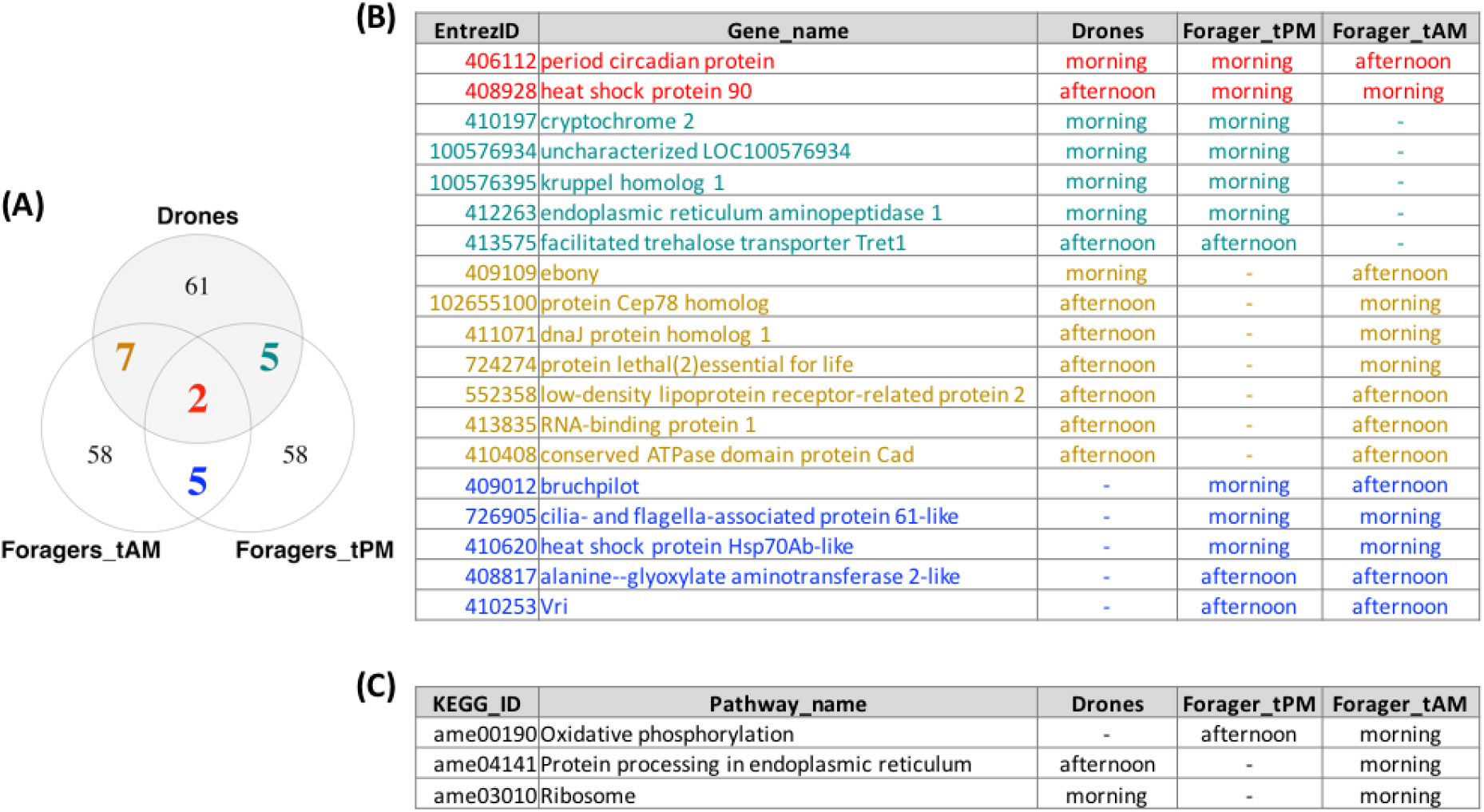
Change in antennal gene expression and signaling pathways with activity. (A) Venn diagram representing number of common and unique genes among drones, afternoon-trained foragers (Foragers_tPM) and morning-trained foragers (Foragers_tAM). (B) All 19 common color coded genes from Venn diagram are listed along with the time of the day when their expression was higher. (C) Significantly enriched genesets (q-value<0.05, GAGE) with the time of the day when their average expression was higher.

We found 5 DEGs (*cry2*, *LOC100576934*, *kruppel homolog1*, *endoplasmic reticulum aminopeptidase1* and *Tret1*) common between drones and afternoon-trained foragers and all of them showed expression changes in the same direction (Fig 6A and 6B). Similar to *per*, expression of *cry2* was higher in the morning in both, drones and afternoon-trained foragers. This further confirms our finding that drones and foragers that are active in the afternoon show similar antennal clock gene rhythms. There were 7 DEGs common between drones and morning-trained foragers out of which 4 (*ebony, protein Cep78 homolog*, *dnaJ protein homolog1* and *protein lethal(2)essential for life*) showed changes in opposite direction suggesting their expression is also associated with the activity rhythm. We also found 5 common DEGs (*bruchpilot*, *cilia- and flagella-associated protein 61-like*, *heat shock protein Hsp70Ab-like*, *alanine--glyoxylate aminotransferase 2-like*, *Vri*) between morning-trained foragers and afternoon-trained foragers. Similar to *per*, *bruchpilot* (*brp*) was higher expressed in the morning in afternoon-trained foragers and higher expressed in the afternoon in morning-trained foragers indicating that the gene is regulated by the activity state. The remaining four genes showed changes in the same direction in both groups appear to be regulated by the time of the day and not the activity state.

In addition to the common genes that showed morning and afternoon expression differences in two or all three experimental groups, we found 61 DEGs that showed changes in the expression only (*p.adj*< 0.05) in the drone antennae (S5 Table). Among these genes were *jun-related antigen* (*Jra*), *Hr38, foraging* (*for*), *dopa decarboxylase* (*Ddc*), *semaphorin-2A*, *calreticullin* (*Crc*), *painless*, *SIFamide receptor* (*SIFR*), *prohormone-2* and many *heat shock proteins* (*hsps*). In the morning-trained foragers, we found 58 DEGs with expression changes between morning and afternoon (e.g. *neurexin 1*, *cwo*, *semaphorin-1A*, *neurobeachin*, *DopEcR* and *SK*) (S5 Table). In the afternoon-trained foragers, there were also 58 DEGs with expression changes between morning and afternoon (e.g. *octopamine receptor* (*Oa1*), *venus kinase receptor* (*Vkr*), *Neural-cadherin* and *Glucose dehydrogenase* (*Gld2*)) (S5 Table).

Geneset enrichment analysis revealed significant enrichment (q.value<0.05) of the following 3 important pathways (Fig 6C and S6 Table) - oxidative phosphorylation (ame00190), protein processing in endoplasmic reticulum (ame04141) and ribosome (ame03010). Oxidative phosphorylation showed significant enrichment only in the foragers and was strongly associated with their activity rhythm. In morning-trained foragers, it was upregulated in the morning; and in afternoon-trained foragers, it was up in the afternoon. Ame04141 was found significantly enriched in drones and morning-trained foragers. It also showed strong association with the activity state. In drones, it was higher in the afternoon (during their mating flight time) while in morning-trained foragers, it was higher in the morning (during their foraging time). Lastly, ame03010 was higher in the morning in drones as well as in morning-trained foragers suggesting no association with the activity state but the time of the day.

## DISCUSSION

In this study, we performed an extensive comparison of gene expression levels between the antennae of honey bee drones and foragers collected at different activity states, i.e. resting vs mating flight activity and resting vs foraging flight activity. The principle findings of these comparisons are: (1.) Drone and worker antennae show sex-specific molecular specialization corresponding to already known morphological and physiological differences (Brockmann and Brückner, 2001; Esslen and Kaissling, 1976). Most obviously, there are only a few olfactory receptor genes with a higher expression in drone antennae, whereas about one third of the annotated olfactory receptor genes are higher expressed in worker antennae. In addition, drone antennae showed a higher expression of genes involved in energy metabolism whereas worker antennae showed a higher expression of genes involved in neuronal communication. (2.) The daily oscillation of the two major clock (*per* and *cry2*) genes in the antennae of drones and foragers correspond to the rest-activity cycles. In foragers, the daily clock gene expression pattern changed with the training time. Drones, which performed mating flights in the afternoon, showed clock gene expression pattern similar to afternoon-trained foragers (Jain and Brockmann, 2018; Sasaki, 1990; Spangler, 1972). (3.) Gene expression comparisons with the time of the day revealed a total of 217 antennal genes regulated by circadian clock and/or onset of activity in honey bees. Out of these only 19 DEGs were common among all 3 groups (drones, morning-trained foragers and afternoon-trained foragers) and remaining 198 DEGs suggested the sex and foraging-time specific regulation of daily antennal gene expression in honey bees.

The most prominent expression differences between drone and worker antennae are highly likely associated with the perception of the queen’s mandibular gland pheromone, which functions as a sex pheromone during mating flights to attract drones (Brockmann et al., 1998; Brockmann et al., 2006). Confirming the findings by Wanner and colleagues (2007), we found the same four olfactory receptor genes, *Or10*, *Or11* (binding 9-ODA, the major mandibular gland component), *Or18*, and *Or170*, and *CEst1* among the top genes showing a drone-biased expression. In addition to this group of genes, we found a very high drone-biased expression for *Shaw*, a gene encoding a potassium channel (Hodge, 2009). To the best of our knowledge there is no report regarding a possible function of *Shaw* in insect olfaction. Further, our gene set enrichment analysis showed that genes involved in oxidative phosphorylation are predominantly higher expressed in drone antennae. Drone antennae have much higher number of sensory neurons than worker antennae and coding in sensory neurons is based on generating action potentials, which are very energy expensive (Attwell and Laughlin, 2001). Similarly, a higher expression of genes involved in protein folding and protein processing might be a consequence of a higher protein turnover rate associated with a higher general activity in drone antennae.

As for the drone antennae, our RNA-seq analysis of the forager antennae clearly corroborated previous microarray studies showing worker-biased gene expression for: *Or63*, *Obp2*, *Obp4*, *Obp11*, *Obp16*, *Obp19*, *Obp21*, *CSP6* and *Cyp6BE1* (Forêt and Maleszka, 2006; Wanner et al., 2007). We also found worker-biased expression of several biogenic amine receptors accompanied by general higher expression of genes involved in the tyrosine/dopamine pathway. Previous studies in honey bees (McQuillan et al., 2012; Vergoz et al., 2009) demonstrated that the expression of dopamine and tyrosine receptors is age and task-dependent and can be modulated by social pheromones. In addition, our GO enrichment analysis suggested higher secondary messenger cascades, cell signaling and extracellular matrix associated genes in drone antennae.

The gene expression differences between drone and worker antennae strongly reflect the different physiological specialization. Drone antennae are specialized to detect small amounts of queen’s pheromone and quickly respond to changes in pheromone concentration (Brockmann et al., 1998; De Bruyne and Baker, 2008). In contrast, forager antennae are predominantly involved in the detection and discrimination of complex odor mixtures which requires “pre-processing” or filtering of the sensory signal sent to the brain. Previous extracellular recordings from the olfactory poreplate sensilla indicated that there might be physiological interactions between the olfactory sensory neurons within one sensillum (Getz and Akers, 1994; Getz and Akers, 1995). Furthermore, inhibitory interactions had been suggested to sharpen and filter the neuronal signal sent to the brain (Andersson et al., 2010; Couto et al., 2005). In addition, the higher expression of genes involved in neural modulation is associated with a high degree of context-dependent plasticity in sensory processing (Bigot et al., 2012; Gadenne et al., 2016; Grosmaitre et al., 2001; McQuillan et al., 2012; Vergoz et al., 2009; Watanabe et al., 2014).

Comparison of gene expression levels in drone and forager antennae between morning and afternoon allowed us to identify olfactory related genes that show temporal changes. Previously, we demonstrated that morning and afternoon feeder time-training phase-shifts the daily oscillation of expression of two major clock genes: *per* and *cry2* in the brain and antennae of honey bee workers (Jain and Brockmann, 2018). Accordingly, *cry2* and *per* showed different expression levels in all the three groups compared (drones, morning-trained foragers, and afternoon-trained foragers). The direction of the change in expression was opposite between morning- and afternoon-trained foragers, and drones showed a similar expression change as the afternoon-trained foragers. Moreover, our qPCR study showed that the daily oscillation of *per* and *cry2* expression in the antenna are similar between drones and afternoon-trained foragers. *Per* and *cry2* expressions peaked during early morning and were the lowest during the late afternoon in both. It has been shown that the clock genes expression rhythm in antennae is necessary for rhythmic olfactory responses and sun compass navigation (Merlin et al., 2009; Tanoue et al., 2004).

As suggested by several earlier studies, the findings of our RNA-seq study confirm that sensory processing in insect antennae appears to be more complex than just detecting odorants and transmitting sensory signals to the brain (Getz and Akers, 1994,1995; Couto et al. 2005; Anderson et al., 2010). Given the sensory specialization of antennae and the relative low number of cell types, whole antenna gene expression analysis provide very robust results (see e.g. our results and those by Wanner et al., 2007). Thus, explorative RNA-seq analysis has the potential to identify molecular players affecting antennal sensory processing as well as to increase our knowledge of possible processing strategies that could be verified by subsequent electrophysiological studies.

## ACKNOWLEDGEMNT

We thank Wolfgang Roessler and Johannes Spaethe for constructive discussion regarding the project and experimental procedure during a research stay of R. Jain at the University of Wuerzburg. We also thank DAAD “A New Passage to India” fellowship for the travel grant to R. Jain.

## AUTHOR CONTRIBUTIONS

R.J. and A.B. designed the experiments. R.J. performed the experiments and analyzed the data. R.J. and A.B. wrote the manuscript.

